# Machine learning-assisted medium optimization revealed the discriminated strategies for improved production of the foreign and native metabolites

**DOI:** 10.1101/2023.02.20.529197

**Authors:** Honoka Aida, Keisuke Uchida, Motoki Nagai, Takamasa Hashizume, Shunsuke Masuo, Naoki Takaya, Bei-Wen Ying

**Affiliations:** School of Life and Environmental Sciences, University of Tsukuba, 1-1-1 Tennodai, Tsukuba, 305-8572 Ibaraki, Japan; Microbiology Research Center for Sustainability, University of Tsukuba, 1-1-1 Tennodai, Tsukuba, 305-8572 Ibaraki, Japan

**Keywords:** production, medium, synthetic pathway, machine learning, bacterial growth, aromatic compound, transcriptome

## Abstract

The medium composition was crucial for achieving the best performance of synthetic construction. What and how medium components determined the production of the synthetic construction remained poorly investigated. To address the questions, a comparative survey with two genetically engineered *Escherichia coli* strains was performed. As a case study, the strains carried the synthetic pathways for producing the aromatic compounds of 4APhe or Tyr, which were common in the upstream but differentiated in the downstream metabolism. Bacterial growth and compound production were examined in hundreds of medium combinations that comprised 48 pure chemicals. The resultant data sets linking the medium composition to bacterial growth and production were subjected to machine learning for improved production. Intriguingly, the primary medium components determining the production of 4PheA and Tyr were differentiated, which were the initial resource (glucose) of the synthetic pathway and the inducer (IPTG) of the synthetic construction, respectively. Fine-tuning of the primary component significantly increased the yields of 4APhe and Tyr, indicating that a single component could be crucial for the performance of synthetic construction. Transcriptome analysis observed the local and global changes in gene expression for improved production of 4APhe and Tyr, respectively, revealing divergent metabolic strategies for producing the foreign and native metabolites. The study demonstrated that ML-assisted medium optimization could provide a novel point of view on how to make the synthetic construction meet the original design.

## Introduction

Designing genetic circuits to construct synthetic metabolic pathways in bacteria is applied for industrial applications and exploring the working principles of living systems [1–4]. In metabolic engineering and synthetic biology, the so-called bottom-up approach using the cellular components as “parts” to rewire the metabolic pathways by genetic reconstruction was used to produce valuable substances in microorganisms [5, 6], e.g., artemisinic acid production [7]. Various tools have been developed to achieve native metabolites or foreign substrates [8], such as the utilization of genome-scale metabolic models (GEMs) [9] and phenotype prediction by machine learning (ML) [10]. Incorporating the synthetic gene circuits into the native regulatory mechanisms was challenging to build the industrially valuable synthetic pathways and create the synthetic metabolism of novel functions [11–13].

In addition to the genetic design, the culture medium was crucial for achieving the best performance of the synthetic construction and modified metabolic pathway [14]. Medium optimization was commonly required to improve the productivity of the endpoint compound (e.g., metabolite, protein) [15–17]. The approaches for medium optimization were intensively developed [18]. The methods based on personal experience and knowledge were popular for quick decisions in laboratories, and those according to the statistic and computational algorithms were used for improved development, e.g., response surface methodology (RSM) [19–21]. Since metabolism was a highly complex network, ML was employed to achieve more significant and efficient optimization [22–25]. Progress in well-established high-throughput technologies and automation primarily benefited the acquisition of large datasets [26, 27], which are essential for ML. The previous studies demonstrated that introducing ML to medium optimization successfully accelerated bacterial growth [28–30] and increased productivity [31, 32].

Despite the successes in genetic design and medium optimization for metabolic engineering, how the optimized medium improved the function of the synthetic pathways (i.e., the production of metabolites) remained undiscovered. What were the decision-making medium components for improved production? The optimized medium was supposed to cause transcriptome reorganization [33, 34], resulting in the refined metabolism benefiting production. How the changes in medium composition contributed to the changes in gene expression of the synthetic pathways? The present study discovered the primary medium components for improved production of synthetic constructions and investigated the changes in gene expression triggered by the fine-tuned medium composition. As a pilot study, two genetically engineered

*Escherichia coli* strains producing two different aromatic compounds, i.e., the native and foreign metabolites, *via* the synthetic pathways of the common upstream and differentiated downstream, were subjected to medium optimization. High-throughput assays were performed to link the medium combinations to bacterial growth and production. The resultant datasets were applied to ML to find the decision-making medium component for improved production. Transcriptome analysis was conducted to discover the metabolic strategy for the increased yields of synthetic constructions.

## Results

### Synthetic pathways for producing aromatic compounds

The genetically engineered *E. coli* strains producing 4-aminophenylalanine (4APhe) or tyrosine (Tyr) were used as a case study. Both strains carried the synthetic pathways, which were in common from glucose to chorismate and differentiated from chorismate to the endpoint metabolites (Fig. 1A). As the wild-type *E. coli* could biosynthesize Tyr but not 4APhe, the compounds of Tyr and 4APhe were designated as the native and foreign metabolites, respectively. Thousands of growth curves were acquired by high-throughput growth assay, and two representative parameters of the growth rate (*r*) and maximal population density (*K*) were calculated in accordance (Fig. 1B). High-performance liquid chromatography (HPLC) was performed to evaluate the production (*P*) of the two compounds at the stationary phase (Fig. 1C). Note that the experimental and analytical approaches used here were all well-established in our previous studies as described in the Materials and Methods.

**Figure 1.**
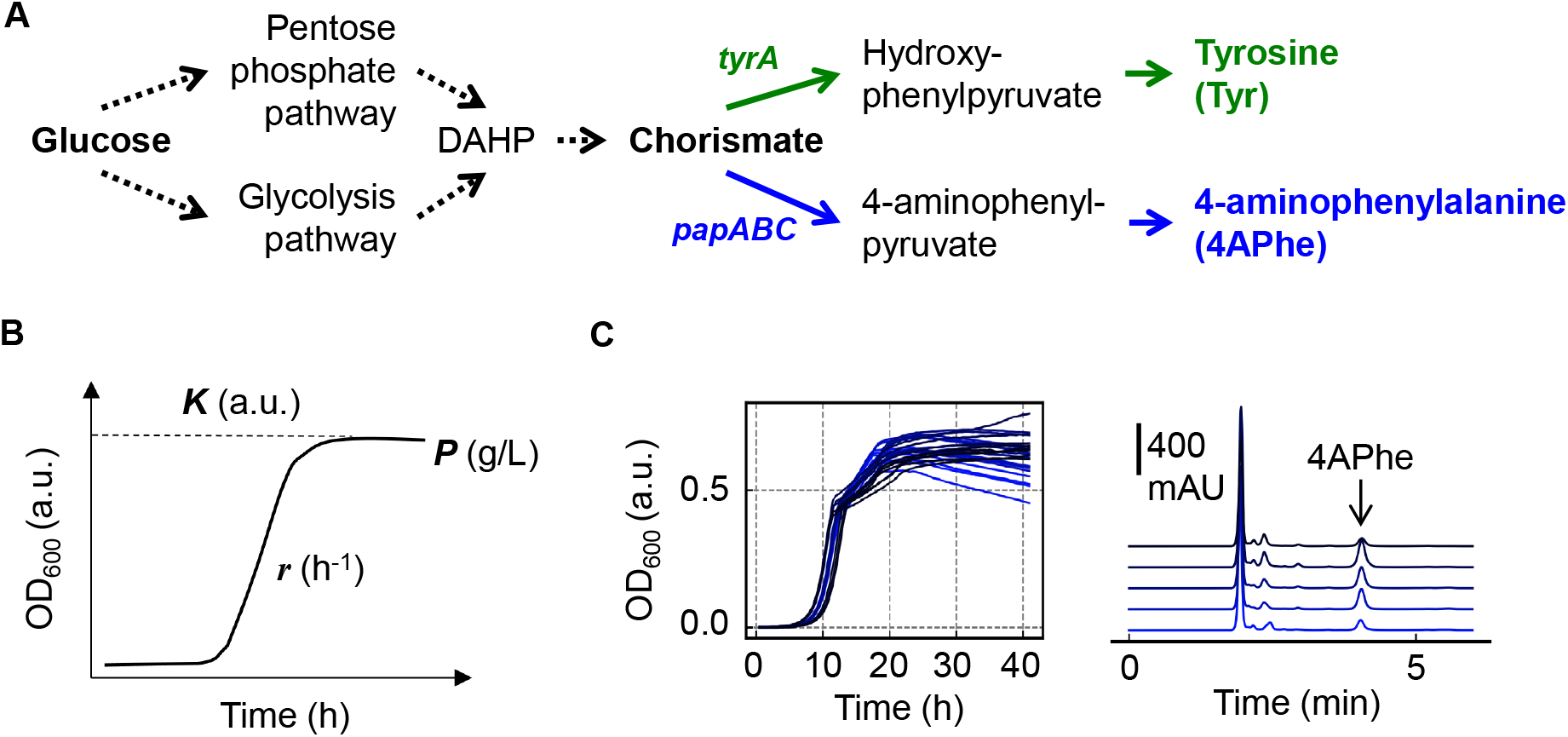
An overview of the experimental and computational analyses. **A.** Flow chart of the synthetic pathways for producing the aromatic compounds. Blue and green indicate the metabolic flows of producing 4APhe and Tyr in two different *E. coli* strains, respectively. **B.** The three parameters used in the study. *K, r*, and *P* represent the maximal density, growth rate, and production yield. The units of the three parameters are indicated. **C.** Examples of the analytical results of bacterial growth and production. The Left and right panels show the growth curves and chromatography peaks in various medium combinations, respectively.

### Linking medium combinations to bacterial growth and production

A total of 48 pure compounds were used to prepare various medium combinations for examining bacterial growth and production, of which 44 were adopted from the previous study using wild-type *E. coli* [28]. Because of the synthetic construction, three antibiotics (chloramphenicol, ampicillin, and streptomycin) and an inducer (IPTG) were added in the present study. 192 and 378 medium combinations were tested for the strains producing 4APhe and Tyr, respectively (Table S1, Table S2). As the pure compounds were ionized in solution, the medium combinations finally consisted of 44 chemical components (Fig. S1, Fig. S2). Repeated assay resulted in thousands of growth curves, and the mean values were used for ML. The final datasets of *r*, *K*, and *P* were 189, 192, and 192 values for 4APhe and 377, 378, and 378 for Tyr. A considerable variation in growth and production was acquired in both strains; however, their distributions were somehow different (Fig. 2A). The distributions of *r*, *K*, and *P* all presented monomodal shape for the strain producing 4APhe (Fig. 2A, upper panels). In comparison, the distribution of *r* was bimodal, although those of *K* and *P* remained monomodal for the strain producing Tyr (Fig. 2A, bottom panels). The dissimilarity indicated that the linkages between growth and production were differentiated in the two synthetic constructions.

**Figure 2.**
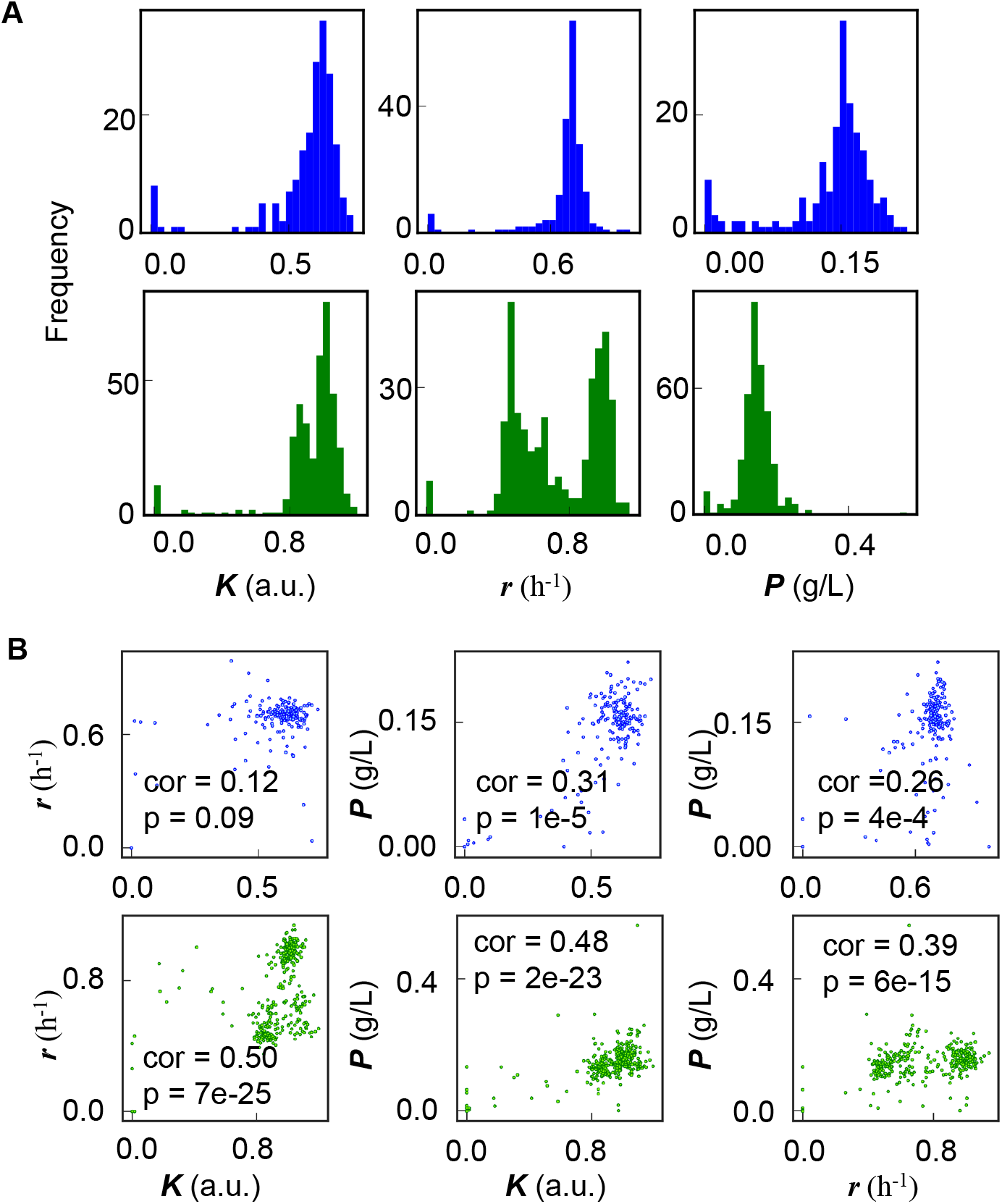
Bacterial growth and production in hundreds of medium combinations. **A.** Profiling of the growth and productivity. Histograms of the maximal density, growth rate, and production yield of the *E. coli* strains grown in various medium combinations are shown from left to right, respectively. The numbers of the tested medium combinations for the *E. coli* strains producing 4APhe and Tyr are 192 and 378, respectively. **B.** Relationships between growth and production. *K*, *r*, and *P* represent the maximal density, growth rate, and production yield. Scatter plots of any pair of *K*, *r*, and *P* are shown. Spearman’s correlation coefficients and the p-values are indicated. Blue and green represent the results of the *E. coli* strains holding the synthetic constructions for producing 4APhe and Tyr, respectively.

The relationship between growth and production was additionally investigated by correlation analysis. The production (*P*) of 4APhe was positively correlated to the growth rate (*r*) and population density (*K*), whereas no correlation was detected between *r* and *K* (Fig. 2B, upper panels). Statistically significant correlations were commonly observed between any pair of *r*, *K*, and *P* in the strain producing Tyr (Fig. 2B, bottom panels), indicating the tight connection between growth and production. The associations among *r*, *K*, and *P* were highly significant in producing the native metabolite, Tyr, and became weaker in producing the foreign one, 4APhe.

### Predicting the primary medium components for production

Gradient-boosted decision tree (GBDT) was applied to predict the decision-making components for growth and production, as described previously [28]. The priority of the medium components (i.e., feature importance) in deciding *r*, *K*, and *P* was evaluated. The results showed that glucose was of high priority in deciding the growth and production (Fig. 3), which was reasonable as glucose was the initial resource for both aromatic compounds (Fig. 1A). The primary components deciding *K* and *r* were glucose and magnesium in the strain producing 4APhe (Fig. 3, upper panels); however, they were both glucose in the strain producing Tyr (Fig. 3, bottom panels). The findings were consistent with the relationship between *r* and *K*, which was no correlation in the strain producing 4APhe and positively correlated in the strain producing Tyr (Fig. 2B, left panels).

**Figure 3.**
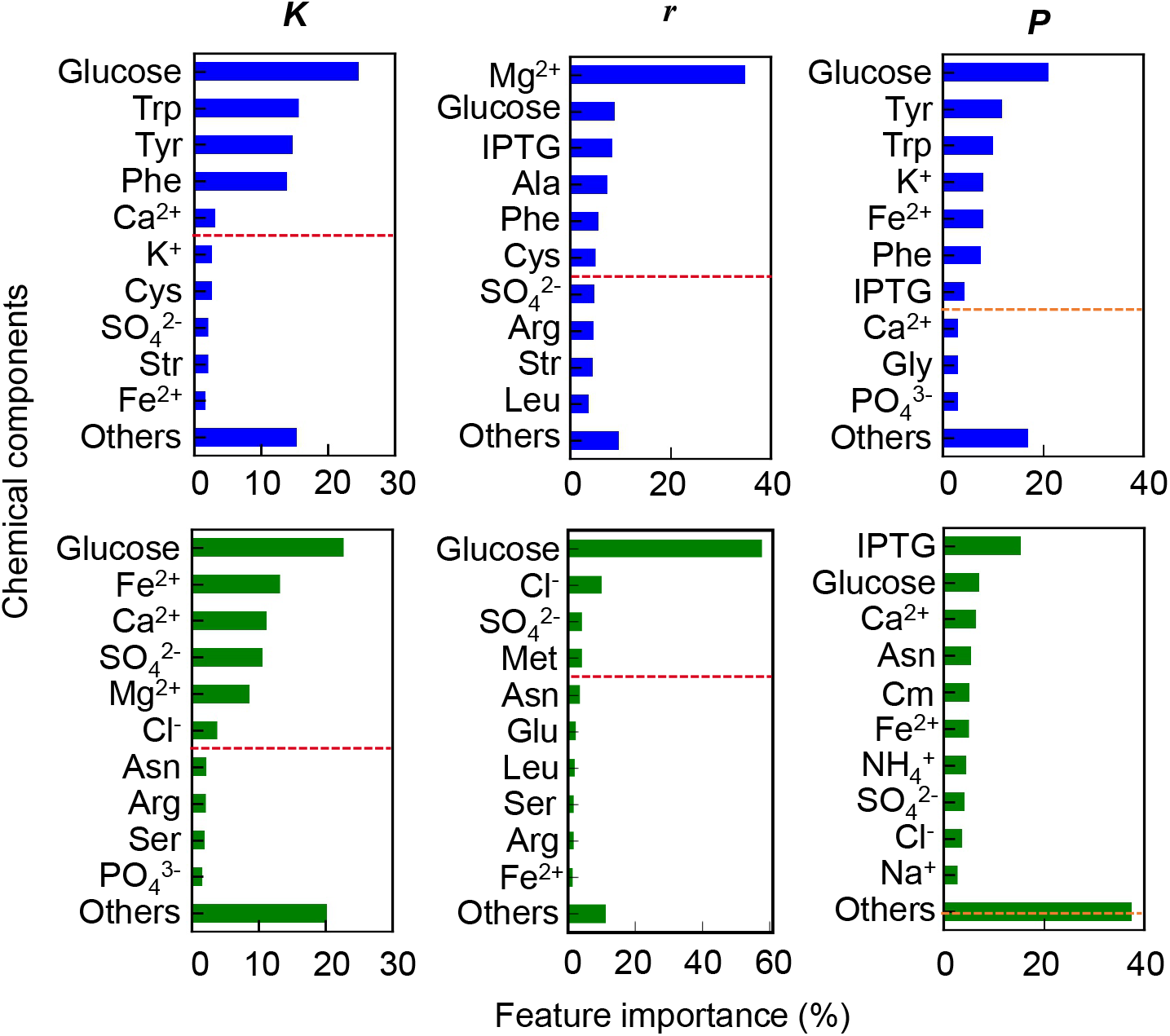
Contribution of the medium components to growth and production. The contribution of each medium component to the maximal density, growth rate, and productivity was predicted by the GBDT model. The top ten primary components affecting the three parameters are displayed, and the rest components are summarized as “Others”. *K*, *r*, and *P* represent the maximal density, growth rate, and production yield. Blue and green represent the results of the *E. coli* strains holding the synthetic pathways for producing 4APhe and Tyr, respectively. The broken lines in orange indicate the border of the cumulated feature importance of ~70%, as summarized in Table S3.

Intriguingly, the decision-making components in the production of 4APhe and Tyr were differentiated: glucose and IPTG, respectively (Fig. 3, right panels). According to the synthetic pathways, glucose was the common initial resource, and IPTG was the common inducer for producing both compounds. Different strategies might be adopted in the two strains to fine-tune the synthetic pathways for production. That is, the metabolic resource-oriented and the transcriptional regulation-dominated strategies in producing 4APhe and Tyr, respectively. In addition, approximately equivalent numbers of the components determined the growth (*r*, *K*) of the two strains (Fig. 3, broken lines, ~70% of feature importance); however, the numbers were significantly different in deciding the productivities of 4APhe and Tyr, i.e., 7 and 14 components, respectively (Fig. 3, Table S3). It revealed that more components participated in the production, not the growth of the strain producing the native metabolite than the foreign one.

### Fine-tuning the concentration of the medium components for improved production

Classification and regression tree (CART) was subsequently applied to estimate the range of chemical concentration, as described previously [30]. The components associated with their concentrations were predicted, and those of high priority in contributing to compound production (*P*) were differentiated in the two strains (Table 1). Glucose and IPTG were the primary components in deciding the production of 4APhe and Tyr, respectively. Trp and Phe were predicted as the decision-making components for producing 4APhe were somehow reasonable, as the synthetic pathways disturbed the native metabolism of tryptophan and phenylalanine biosynthesis [35]. In comparison, the two amino acids insignificantly contributed to the production of Tyr, although their biosynthesis was interrupted by the synthetic pathways. The differentiation in the primary components for substrate production in the two strains was highly consistent with that predicted with GBDT, despite a few dissimilarities between the two ML models.

**Table 1.** Chemical concentrations and productivity predicted by CART. The components of high priority in contributing to productivity (*P*) are shown. The predicted concentrations of the components for improved productivity are indicated.

|  | 4APhe | Tyr |
| --- | --- | --- |
|  | Trp > 0.06 |  |
| Concentration (mM) | **Glucose > 6.3** | **0.01 < IPTG < 0.03** |
| | $\text{Cu}^{2+} > 1.0 \times 10^{-5}$ | Aminobenzoic acid > 0.001 |
|  | Phe > 0.02 |  |
| *P* (g/L) | 0.15 | 0.56 |

Medium optimization by fine-tuning the concentrations of glucose and IPTG was performed to demonstrate whether the predicted chemical concentrations improved the production. The results showed that the yields of both compounds were changed in response to the concentration gradients of the two primary components (Fig. 4A). The chemical concentrations presenting high yields were precisely within the predicted ranges. The concentrations of glucose and IPTG for the best production of 4APhe and Tyr were finally decided as 40 and 0.025 mM, respectively. Additional experiments verified the yields of 4APhe and Tyr were 0.30 and 0.65 g/L, which were both higher than (or equivalent to) the prediction of 0.15 and 0.56 g/L, respectively (Table 1).

**Figure 4.**
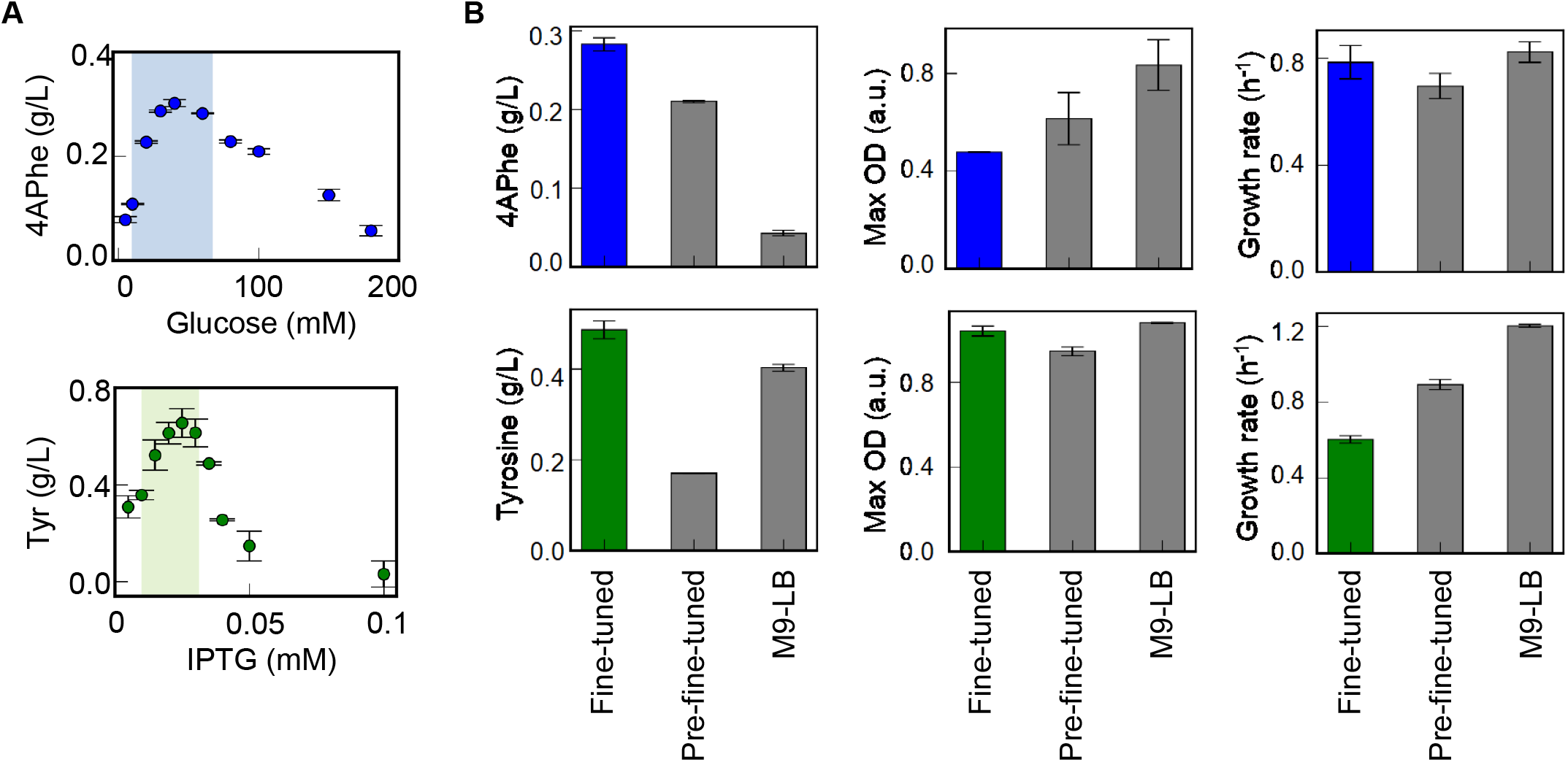
Changes in production in response to medium variation. **A.** Testing the primary components for production. The upper and bottom panels indicated the production of 4APhe and Tyr, respectively. Ten medium combinations of identical concentrations of 47 components but varying concentrations of glucose or IPTG were tested. The shadows represent the concentration gradients of the primary components (glucose or IPTG) predicted by the CART model. Standard errors of experimental replications are indicated. **B.** Testing the representative media. The media of fine-tuned and pre-fine-tuned differed in the concentration of the primary component, i.e., glucose for 4APhe and IPTG for Tyr. M9-LB represents the mixture of the M9 and LB media used previously. Standard errors of experimental replications are indicated.

Furthermore, whether the fine-tuning of the primary components improved the production in the culture of a larger volume was examined. A scaleup to 5 mL of culture was conducted to compare the production of 4APhe and Tyr in three different media, i.e., the fine-tuned, pre-fine-tuned, and M9-LB media. The highest yields of both 4APhe and Tyr were acquired in the fine-tuned media (Fig. 4B, left panels), comparable to those cultured in 200 μL (Fig. 4A). Intriguingly, the increased yields linked to the lower growth rates and maximal population densities (Fig. 4B, right two panels). It strongly suggested that the trade-off between growth and production should be considered in synthetic constructions.

### Changes in gene expression caused by the fine-tuned medium composition

Transcriptome analysis was performed to investigate whether and how a single fine-tuned medium component triggered the changes in gene expression. Differential expression genes (DEGs) mediated by medium optimization were identified more in the stationary phase than in the exponentially growing phase (Fig. 5A). Only a few DEGs overlapped between the exponential and the stationary phases. Significant changes in gene expression commonly occurred in the phase for production but not growth, independent of the variation in the fine-tuned components (i.e., glucose and IPTG) and final products (i.e., 4APhe and Tyr). It indicated that ML-assisted prediction was practical for finding the critical component for efficient medium optimization, and tuning a single medium component was sufficient to improve the performance of the synthetic construction.

**Figure 5.**
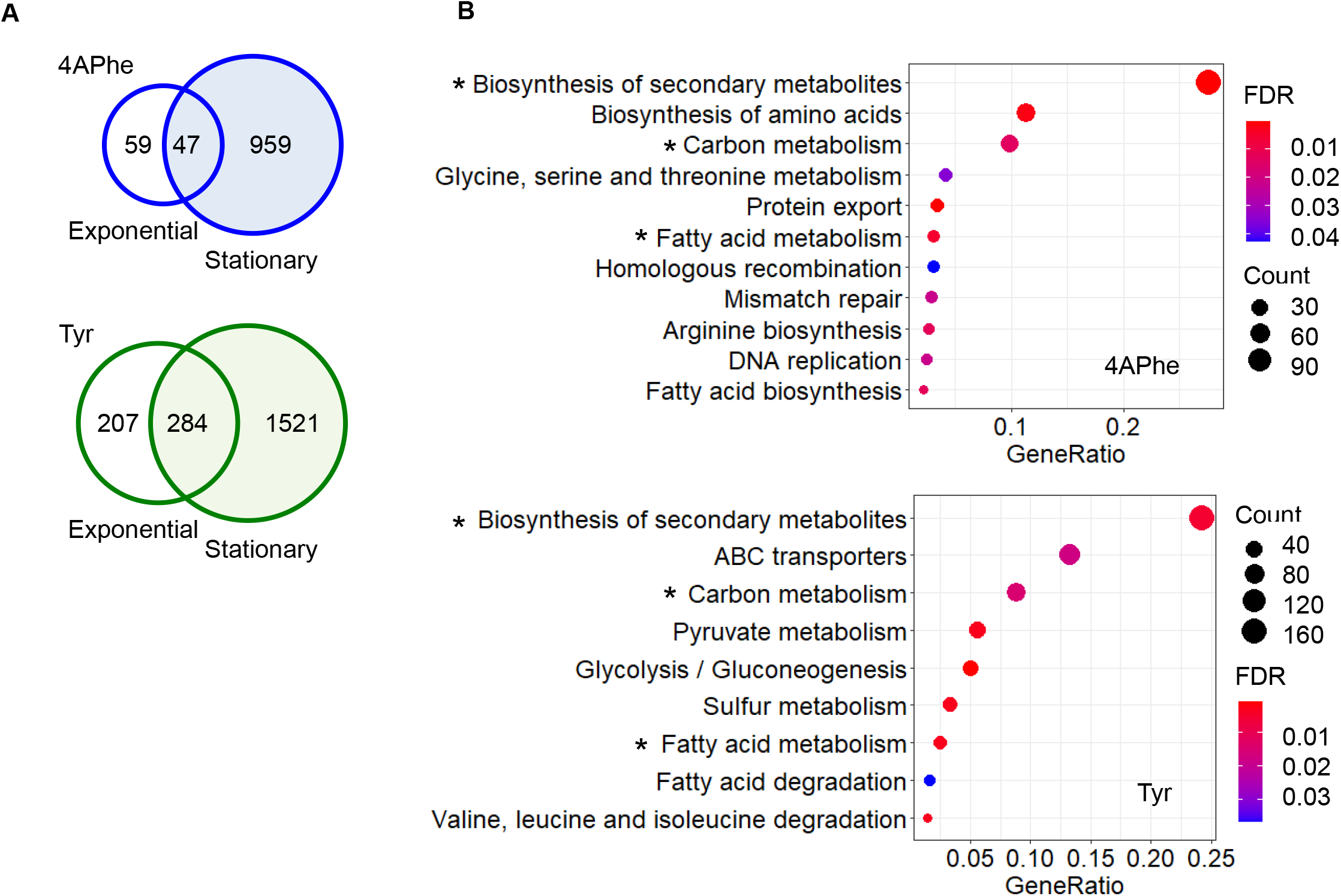
Differentiated gene expression in response to medium optimization. **A.** Numbers of DEGs in response to medium optimization. Open and shadowed circles stand for the exponential and stationary phases, respectively. Blue and green represent the results of the *E. coli* strains holding the synthetic pathways for producing 4APhe and Tyr, respectively. **B.** Enriched metabolic pathways in DEGs. The KEGG pathways significantly enriched in the DEGs are indicated (FDR < 0.05). The upper and bottom panels indicate the strains producing 4APhe and Tyr, respectively. Gene ratio is the ratio of DEGs to all genes annotated in the pathway. Color gradation and the size of circles indicate the statistical significance represented by FDR and the number of DEGs annotated in the pathway.

Functional enrichment analysis of DEGs showed that a total of 11 and 9 KEGG pathways significantly (FDR < 0.05) fluctuated in the strains producing 4APhe and Tyr, respectively (Fig. 5B). However, the numbers of DEGs varied between the two strains, i.e., 1,006 and 1,805 DEGs. Three native metabolic pathways (biosynthesis of secondary metabolites, carbon metabolism, and fatty acid metabolism) were commonly enriched in response to the medium optimization (Fig. 5B, asterisks), which might be due to the universal upstream of the synthetic pathways for the production of 4APhe and Tyr.

In addition, the improved production of 4APhe was mainly attributed to the transcriptional regulation of downstream pathways close to the endpoint metabolite (Fig. 6A, blue). In comparison, the improved production of Tyr was accompanied by transcriptional changes across entire pathways from the initial resource to the endpoint metabolite (Fig. 6A, green). Despite the universal upstream metabolism, the expression changes of the genes that participated in the synthetic pathways occurred at the local and global scale for 4APhe and Tyr, respectively. It indicated that the transcriptome reorganization for improved production was differentiated between the foreign and native metabolites. Whether the fine-tuned component was a resource initiating the metabolism or an inducer regulating the gene expression must be crucial for metabolic strategy (Fig. 6B).

**Figure 6.**
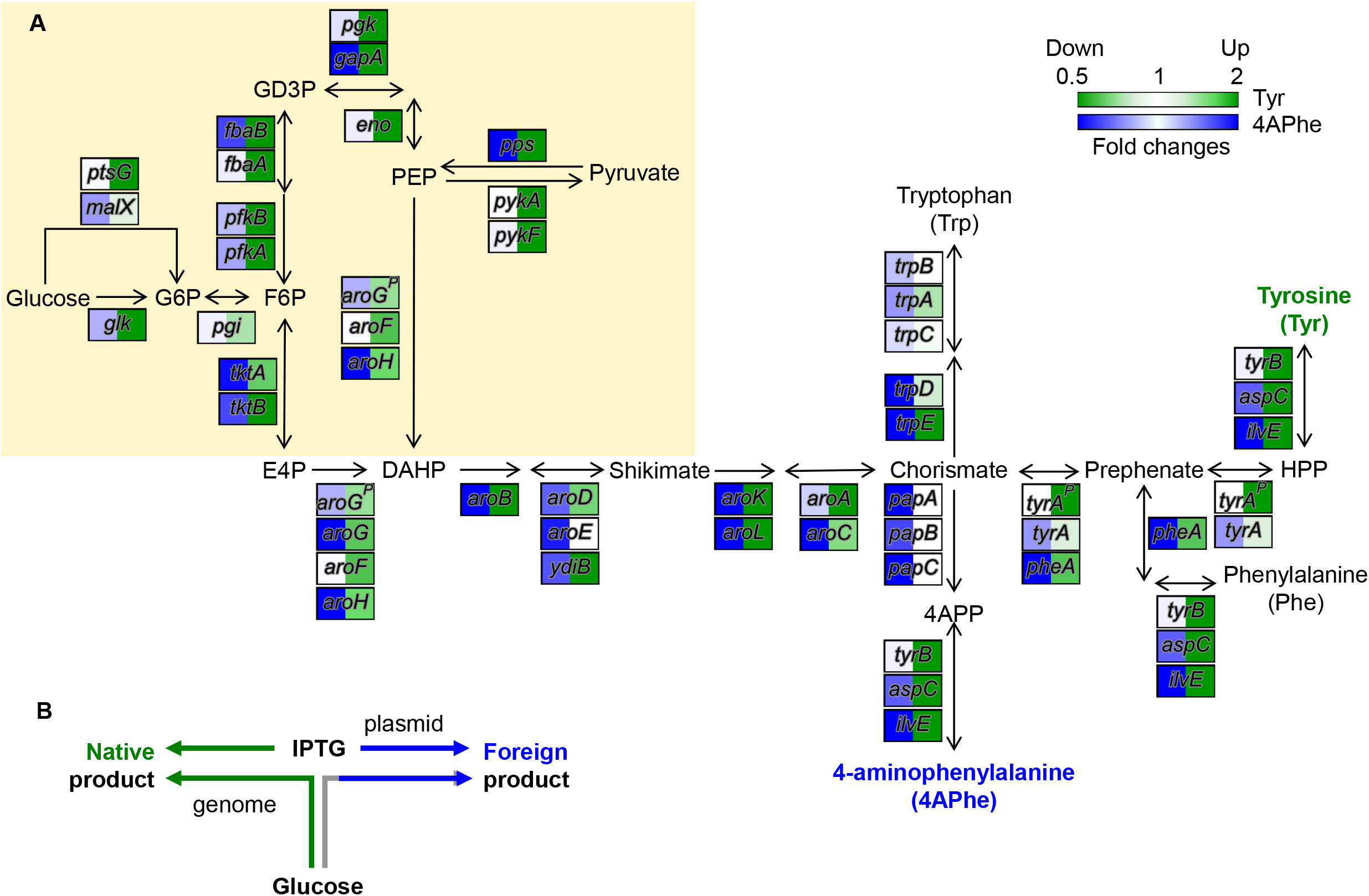
Transcriptional changes in the synthetic pathways. **A.** Transcriptional changes of the genes participated in the metabolic pathways for production. The genes and chemical compounds participating in the metabolic pathways are shown. Color gradation in blue and green represent the fold changes of gene expression caused by the medium optimization. Shadow in light yellow indicates the upstream glycolysis and pentose phosphate pathways in common for producing 4APhe and Tyr. **B.** Schematic drawing of the working principle of synthetic pathways.

## Discussion

Combining the data-driven approach and the transcriptome analysis allowed us to find the different strategies of metabolic optimization between foreign and native metabolites. The highest priority chemicals contributing to the production of 4APhe and Tyr were glucose and IPTG, respectively. The genes for aromatic amino acids biosynthesis and transcriptional feedback repression (i. e., *tyrA, trpE, pheA*, and *tyrR*) were mutated or deleted in the strains producing 4APhe [35, 36]. The feedback repression was released so that the limiting factor for 4APhe production became the initial resource, glucose. On the other hand, as the feedback mechanism remained regular in the strain producing Tyr, the IPTG-induced expression level of the enzyme (i.e., *tyrA*) might play an essential role.

Fine-tuning of a single medium component caused the transcriptome reorganization of the overall pathways for Tyr production, indicating that balancing metabolism was essential for producing the native metabolite. Since the native proteins and metabolites participate in the entire cellular process, their overall activation might cause metabolic imbalance and be lethal. The finding supported that the overexpression of native proteins caused severe inhibition of *E. coli* growth [37]. It might be why the inducer IPTG was predicted to be the primary component for producing Tyr. The appropriate gene expression level for a balanced metabolism must have been the key to native production (metabolite). In addition, only the local but not global reorganization of the transcriptome occurred in producing the foreign compound, 4APhe. The local metabolic modification might be sufficient for improved production because of the fewer interactions of the foreign metabolite to the native pathways. The finding supported that metabolic imbalance occurred once the native but not the orthologous dihydrofolate reductase (DHFR) was overexpressed [38].

Adding the inducer at the beginning of cell culture for production seemed highly applicable. Although the inducer IPTG was present from the beginning of the culture, much more DEGs were detected in the stationary phase than in the exponential phase. It suggested that the proper concentration of IPTG at the initial growth phase could affect the transcriptome for better production in the stationary phase. Adding IPTG in the medium adjusted the cost of growth and protein production in *E. coli* [39]. Nevertheless, the IPTG induction at the inoculum did not cause a growth burden but allowed high penicillin productivity in *E. coli* [40], which supported the present finding. The traditional protocol of IPTG induction at the stationary phase was somehow nonessential. In addition, the optimal IPTG concentration for Tyr production (Fig. 4B) seemed to be ~ 10-fold lower than those generally used for recombinant protein production [41, 42]. The improved production of the synthetic construction was not simply due to the induced expression of the critical enzyme but the transcriptional reorganization of the whole metabolic pathway (Fig. 6).

Whether the primary component changed in the culture medium of a larger volume was tested. The production of 4APhe was highly responsive to glucose in a 5 mL culture (Fig. S4) within the concentration gradient as in the 200 μL culture (Fig. 4A). It demonstrated that glucose was the decision-making component in a large-volume culture. In addition, the fine-tuned medium was highly competitive for producing 4APhe compared to the commercially available rich media (Fig. S5). Despite the highest yield in the fine-tuned medium, the growth rate and maximal population density were lower than those in the commercial media (Fig. S5). The trade-off between production and growth demonstrated that the fine-tuned medium was directed explicitly toward production but not growth.

Although the tested medium combinations were of limited variety, the high-throughput assay combined with ML-assisted medium optimization was practical. As the increased richness of the media did not always result in improved growth and production (Fig. S3), the nutritional balance of media was supposed to be critical. So far, finding an ideal medium for the synthetic construction to reach its best performance remains challenging because both the medium and the metabolism are highly complex. As a case study, the present study provided a pilot trial of medium optimization with the decision-making component for the synthetic pathways.

The fine-tuned media led to the reorganized transcriptomes for improved production, regardless of the foreign or native metabolites. The differentiation in transcriptome reorganization revealed the different metabolic strategies for producing foreign and native compounds. Simply increasing the initial resource (e.g., glucose) might lead to an increased amount of the foreign product, e.g., 4APhe. It might be because of the weak interaction between the foreign product and the native metabolism. However, a balanced induction of the whole pathway was required for increasing the amount of the native product, e.g., Tyr. It might be caused by the tight regulation affecting the overall metabolic balance. The ML-assisted high-throughput approaches benefited novel findings in transcriptional and metabolic principles for constructing synthetic pathways, which was valuable for using the bacterial cell as a factory.

## Materials and Methods

### E. coli strains

Two genetically reconstructed *E. coli* strains that produced two different aromatic compounds from chorismate via the shikimic acid pathway were used. The *E. coli* strain *NST37(DE3)/ΔpheLA* harboring pET-pfpapA/pCDF-pfpapBC/pACYC-aroG4 was previously constructed to produce 4-aminophenylalanine (4APhe) production [35]. The expression of *papABC* mediated the production of 4APhe. The *E. coli* strain BL21(DE3) harboring pET-tyrA/pACYC-aroG^fbr^ was constructed to produce tyrosine (Tyr), as described previously [43]. Both products were induced by isopropyl β-D-1-thiogalactopyranoside (IPTG) (Fig. 1A).

### Natural media

The media of Instant LB, APS, Plusgrow II, and Lennox were obtained commercially (BD Biosciences), and M9LB was prepared in the lab. The M9LB medium comprises 24 g/L Na_2_HPO_4_, 12 g/L KH_2_PO_4_, 1 g/L NH_4_Cl, 0.5 g/L NaCl, 0.05 g/L thiamine HCl, 0.5 g/L MgSO_4_-7H_2_O, 0.015 g/L CaCl_2_-H_2_O, 5g/L Yeast extract, 10g/L Tryptone and 2mL/L Hunter’s trace elements, which contain 22 g/L ZnSO_4_-H_2_O, 11g/L H_3_BO_3_, 5 g/L MnCl_2_-4H_2_O, 5 g/L FeSO_4_-7H_2_O, 1.6 g/L CoCl_2_-6H_2_O and CuSO_4_/5H_2_O. The chemical compounds are all commercially available (Wako).

### Preparation of medium combinations

A total of 48 pure compounds were used, of which 44 were previously determined [28], and four were newly added. The four additives were the inducer (i.e., IPTG) and antibiotics (i.e., ampicillin, chloramphenicol, and streptomycin). The pure compounds were all purchased commercially (Wako). The concentration gradients of these compounds were determined as the same as in the previous study [28]. In brief, the minimal and maximal concentrations were zero and 10~100 folds of the concentrations present in the commonly used media, respectively. The chemical concentrations were varied on a logarithmic scale. The stock solutions of these 48 compounds were prepared and stored in small aliquots (100~1,000 μL) at −30 °C, as described previously. The stock solutions were used only once, and the remainder was discarded to avoid deterioration of compound quality due to repeated thawing and freezing. The medium combinations were prepared by mixing these stock solutions before the growth assay. Finally, 192 and 378 medium combinations were tested in the growth assays for the *E. coli* strains that produce 4APhe and Tyr, respectively (Table S1).

### Bacterial growth assay

The cell stocks of exponentially growing *E. coli* cells (OD_600_ ≈ 0.01-0.1) were prepared before the growth assay for repeated use, as described previously [44]. The cell stocks (cell culture aliquots) were used only once, and the remaining culture was discarded. The high-throughput growth assays were performed using the 96-well microplates (Costar) and the plate reader (Epoch2, BioTek), as described in the previous studies [28]. Every four to six wells (200 μl per well) in different positions were tested for each medium combination. The temporal changes of the *E. coli* cells growing at 37°C were recorded by reading absorbance at 600 nm for 30~48 h in 30-min intervals. In addition, the growth assay in a test tube was performed with 5 ml of the culture at 37 °C and 200 rpm using a bioshaker (Taitec). The cell culture was temporally sampled by taking 0.1~1.0 mL of culture to measure OD600 (Beckman DU730), as previously described in detail [45].

### Growth data processing and calculation

The temporal OD_600_ reads were exported from the plate reader and processed with Python. The growth parameters *r* and *K* were evaluated according to previous reports [30, 46] using a previously developed Python program [30]. In brief, *r* was defined as the mean of three continuous logarithmic slopes of every two neighboring OD_600_ values within the exponential growth phase using “gradient” in the “NumPy” library. *K* was calculated as the mean of three continuous OD_600_ values, including the maximum, determined using “argmax” in the “NumPy” library.

### High-performance liquid chromatography

The amounts of 4APhe and Tyr produced by the *E. coli* cells grown in various media were evaluated by high-performance liquid chromatography (HPLC). The cell cultures were centrifuged at 16,000 rpm for 5 min (Model 3700, Kubota), and the supernatants were collected and subjected to HPLC (1260 Infinity, Agilent Technologies). The flow rate of HPLC was set at 0.8 mL/min. A HiQ sil C18HS-3 column (Siri Instrument) was equipped to detect 4APhe. The initial mobile phase was 20 mM KH_2_PO_4_ (pH 7.0) and pure methanol (Sigma-Aldrich) in the ratio of 98 to 2 and maintained for 7 min. The methanol concentration was gradually increased to 50% in 5 minutes and then maintained for 5 minutes. Subsequently, the ratio of methanol was gradually reduced to 2% for 2 min, and the flow was kept for 4 min. In addition, a TSKgel ODS-100V column (Tosoh) was used to detect Tyr. The initial mobile phase was 10 mM ammonium formate (pH 7.0) and pure acetonitrile (Merck Millipore) in the ratio of 95 to 5 and maintained for 8 minutes. The acetonitrile concentration was increased to 50% in 6 min and then maintained for 2 min. Subsequently, the ratio of acetonitrile was gradually reduced to 5% in 4 min, and the flow was kept for 5 min. *p-*Amino-L-phenylalanine (Sigma-Aldrich) and L-tyrosine (Wako) were used as the standards of 4APhe and Tyr, respectively. The amounts (productivity, *P*) of 4APhe and Tyr were calculated according to the peak areas at the corresponding retention times.

### Machine learning

Machine learning was performed with Python, as described previously[28, 30]. Two machine learning models of gradient-boosted decision tree (GBDT) and classification and regression trees (CART) were applied to the datasets that linked the parameters of growth (i.e., *r* and *K*) and productivity (i.e., *P*) to the medium combinations. The chemical concentrations of the medium combinations were transformed into logarithmic values in the datasets. “GradientBoostingRegressor” in the “ensemble” module and “DecisionTreeRegressor” in the “tree” module were used for GBDT and CART, respectively. Both were in the “scikit-learn” library. A fivefold nested cross-validation was performed to evaluate the GBDT model. A grid search was used for the hyperparameter search in the GBDT model, with “n_estimators” set to 300. The values of “learning_rate” and “max_depth” were searched in the ranges of 0.005 ~ 0.5 in increments of 0.005 and 1 ~ 4 in increments of 1, respectively. All other hyperparameters were used as default. The “feature_importance_” values were calculated by fivefold cross-validation. The mean value of five repeated calculations was used as the GBDT predicted output. In the CART model, “criterion” was set to “mse”, “max_depth” was set to 5, and all other hyperparameters were used as default.

### RNA purification and RNA sequencing

The *E. coli* cells were cultured in 5 ml of the fresh media and rotated at 200 rpm at 37□. Independent cultures as biological replicates of transcriptomes were performed. The cell cultures of both exponential and stationary growth phases were collected, as described previously [47, 48]. The total RNAs were purified using RNeasy Mini Kit (QIAGEN) and RNase-Free DNase Set (QIAGEN) according to the product instructions. The eluted RNAs were suspended with RNase-free water and subsequently subjected to RNAseq. The rRNAs were removed using Ribo-Zero Plus rRNA Depletion Kit (Illumina), and mRNA preparation was performed with Ultra Directional RNA Library Prep Kit for Illumina (NEBNext). The paired-end sequencing (150 bp × 2) was performed with Novaseq6000 (Illumina) by Chemical Dojin Co. Ltd.

### Transcriptome data processing and analysis

The computational and statistical analyses were all performed using R [49]. The reference genome sequences of *E. coli* W3110 and BL21(DE3) were obtained from the GenBank of the accession numbers ASM1024v1 and CP001509.3, respectively. The trimmed reads were mapped to the reference genome sequences with Bowtie2 [50]. The transcriptome datasets were deposited in the DNA Data Bank of Japan (DDBJ) under the accession number DRA013628. The differential expressed genes (DEGs) were determined using DESeq2 [51] package, with a criterion of FDR < 0.05 [52], where the read counts were directly used. The enrichment analysis was performed using the globally normalized datasets, as previously described [47, 53]. KEGG pathways [54, 55] were enriched using “enrichKEGG” in clusterProfiler [56], where “pAdjustMethod” was set to “fdr,” and “organism” was set to “ecj” and “ebe” for the *E. coli* strains that produced 4APhe and Tyr, respectively.

## Acknowledgments

This work was partially supported by the JSPS KAKENHI Grant-in-Aid for Scientific Research (B) (19H03215 to BWY) and Grant-in-Aid for Challenging Exploratory Research (21K19815 to BWY).

## Competing interests

The authors (BWY and HA) have competing interests. The medium combinations were related to a patent under the control number 2021-171528 (Japan).

## References

1. Otero-Muras I, Carbonell P: Automated engineering of synthetic metabolic pathways for efficient biomanufacturing. Metab Eng 2021, 63:61–80.

2. Long B, Fischer B, Zeng Y, Amerigian Z, Li Q, Bryant H, Li M, Dai SY, Yuan JS: Machine learning-informed and synthetic biology-enabled semi-continuous algal cultivation to unleash renewable fuel productivity. Nat Commun 2022, 13:541.

3. Fong SS: Computational approaches to metabolic engineering utilizing systems biology and synthetic biology. Comput Struct Biotechnol J 2014, 11:28–34.

4. Jouhten P: Metabolic modelling in the development of cell factories by synthetic biology. Comput Struct Biotechnol J 2012, 3:e201210009.

5. Stephanopoulos G: Synthetic Biology and Metabolic Engineering. ACS Synthetic Biology 2012, 1:514–525.

6. Cameron DE, Bashor CJ, Collins JJ: A brief history of synthetic biology. Nat Rev Microbiol 2014, 12:381–390.

7. Ro DK, Paradise EM, Ouellet M, Fisher KJ, Newman KL, Ndungu JM, Ho KA, Eachus RA, Ham TS, Kirby J, et al: Production of the antimalarial drug precursor artemisinic acid in engineered yeast. Nature 2006, 440:940–943.

8. Keasling JD: Synthetic biology and the development of tools for metabolic engineering. Metab Eng 2012, 14:189–195.

9. Gu C, Kim GB, Kim WJ, Kim HU, Lee SY: Current status and applications of genome-scale metabolic models. Genome Biol 2019, 20:121.

10. Carbonell P, Radivojevic T, García Martín H: Opportunities at the Intersection of Synthetic Biology, Machine Learning, and Automation. ACS Synth Biol 2019, 8:1474–1477.

11. Satowa D, Fujiwara R, Uchio S, Nakano M, Otomo C, Hirata Y, Matsumoto T, Noda S, Tanaka T, Kondo A: Metabolic engineering of E. coli for improving mevalonate production to promote NADPH regeneration and enhance acetyl-CoA supply. Biotechnol Bioeng 2020, 117:2153–2164.

12. Huccetogullari D, Luo ZW, Lee SY: Metabolic engineering of microorganisms for production of aromatic compounds. Microb Cell Fact 2019, 18:41.

13. Larroude M, Celinska E, Back A, Thomas S, Nicaud JM, Ledesma-Amaro R: A synthetic biology approach to transform Yarrowia lipolytica into a competitive biotechnological producer of β-carotene. Biotechnol Bioeng 2018, 115:464–472.

14. Overmann J, Abt B, Sikorski J: Present and Future of Culturing Bacteria. Annu Rev Microbiol 2017, 71:711–730.

15. Abuhena M, Al-Rashid J, Azim MF, Khan MNM, Kabir MG, Barman NC, Rasul NM, Akter S, Huq MA: Optimization of industrial (3000 L) production of Bacillus subtilis CW-S and its novel application for minituber and industrial-grade potato cultivation. Sci Rep 2022, 12:11153.

16. Krause M, Neubauer A, Neubauer P: The fed-batch principle for the molecular biology lab: controlled nutrient diets in ready-made media improve production of recombinant proteins in Escherichia coli. Microb Cell Fact 2016, 15:110.

17. Choi GH, Lee NK, Paik HD: Optimization of Medium Composition for Biomass Production of Lactobacillus plantarum 200655 Using Response Surface Methodology. J Microbiol Biotechnol 2021, 31:717–725.

18. Singh V, Haque S, Niwas R, Srivastava A, Pasupuleti M, Tripathi CK: Strategies for Fermentation Medium Optimization: An In-Depth Review. Front Microbiol 2016, 7:2087.

19. Aguirre AM, Bassi A: Investigation of biomass concentration, lipid production, and cellulose content in Chlorella vulgaris cultures using response surface methodology. Biotechnol Bioeng 2013, 110:2114–2122.

20. Bezerra MA, Santelli RE, Oliveira EP, Villar LS, Escaleira LA: Response surface methodology (RSM) as a tool for optimization in analytical chemistry. Talanta 2008, 76:965–977.

21. Latha S, Sivaranjani G, Dhanasekaran D: Response surface methodology: A non-conventional statistical tool to maximize the throughput of Streptomyces species biomass and their bioactive metabolites. Crit Rev Microbiol 2017, 43:567–582.

22. Packiam KAR, Ooi CW, Li F, Mei S, Tey BT, Ong HF, Song J, Ramanan RN: PERISCOPE-Opt: Machine learning-based prediction of optimal fermentation conditions and yields of recombinant periplasmic protein expressed in Escherichia coli. Comput Struct Biotechnol J 2022, 20:2909–2920.

23. Lawson CE, Martí JM, Radivojevic T, Jonnalagadda SVR, Gentz R, Hillson NJ, Peisert S, Kim J, Simmons BA, Petzold CJ, et al: Machine learning for metabolic engineering: A review. Metabolic Engineering 2021, 63:34–60.

24. Kim GB, Kim WJ, Kim HU, Lee SY: Machine learning applications in systems metabolic engineering. Curr Opin Biotechnol 2020, 64:1–9.

25. Cuperlovic-Culf M: Machine Learning Methods for Analysis of Metabolic Data and Metabolic Pathway Modeling. Metabolites 2018, 8.

26. Gilpin W, Huang Y, Forger DB: Learning dynamics from large biological data sets: Machine learning meets systems biology. Current Opinion in Systems Biology 2020, 22:1–7.

27. Suthers PF, Foster CJ, Sarkar D, Wang L, Maranas CD: Recent advances in constraint and machine learning-based metabolic modeling by leveraging stoichiometric balances, thermodynamic feasibility and kinetic law formalisms. Metabolic Engineering 2021, 63:13–33.

28. Aida H, Hashizume T, Ashino K, Ying BW: Machine learning-assisted discovery of growth decision elements by relating bacterial population dynamics to environmental diversity. Elife 2022, 11.

29. Hiura S, Koseki S, Koyama K: Prediction of population behavior of Listeria monocytogenes in food using machine learning and a microbial growth and survival database. Sci Rep 2021, 11:10613.

30. Ashino K, Sugano K, Amagasa T, Ying BW: Predicting the decision making chemicals used for bacterial growth. Sci Rep 2019, 9:7251.

31. Kumar P, Adamczyk PA, Zhang X, Andrade RB, Romero PA, Ramanathan P, Reed JL: Active and machine learning-based approaches to rapidly enhance microbial chemical production. Metab Eng 2021, 67:216–226.

32. Zheng Z-Y, Guo X-N, Zhu K-X, Peng W, Zhou H-M: Artificial neural network – Genetic algorithm to optimize wheat germ fermentation condition: Application to the production of two anti-tumor benzoquinones. Food Chemistry 2017, 227:264–270.

33. Feugeas JP, Tourret J, Launay A, Bouvet O, Hoede C, Denamur E, Tenaillon O: Links between Transcription, Environmental Adaptation and Gene Variability in Escherichia coli: Correlations between Gene Expression and Gene Variability Reflect Growth Efficiencies. Mol Biol Evol 2016.

34. Blair JMA, Richmond GE, Bailey AM, Ivens A, Piddock LJV: Choice of Bacterial Growth Medium Alters the Transcriptome and Phenotype of Salmonella enterica Serovar Typhimurium. PLOS ONE 2013, 8:e63912.

35. Masuo S, Zhou S, Kaneko T, Takaya N: Bacterial fermentation platform for producing artificial aromatic amines. Sci Rep 2016, 6:25764.

36. Tribe DE: Novel microorganism and method. United States: Austgen Biojet International Pty Ltd; 1987.

37. Kitagawa M, Ara T, Arifuzzaman M, Ioka-Nakamichi T, Inamoto E, Toyonaga H, Mori H: Complete set of ORF clones of Escherichia coli ASKA library (A Complete S et of E. coli K −12 ORF A rchive): Unique Resources for Biological Research. DNA Research 2005, 12:291–299.

38. Bhattacharyya S, Bershtein S, Yan J, Argun T, Gilson AI, Trauger SA, Shakhnovich EI: Transient protein-protein interactions perturb E. coli metabolome and cause gene dosage toxicity. eLife 2016, 5:e20309.

39. Malakar P, Venkatesh KV: Effect of substrate and IPTG concentrations on the burden to growth of Escherichia coli on glycerol due to the expression of Lac proteins. Applied Microbiology and Biotechnology 2012, 93:2543–2549.

40. Ramírez OT, Zamora R, Espinosa G, Merino E, Bolívar F, Quintero R: Kinetic study of penicillin acylase production by recombinant E. coli in batch cultures. Process Biochemistry 1994, 29:197–206.

41. Lipničanová S, Legerská B, Chmelová D, Ondrejovič M, Miertuš S: Optimization of an Inclusion Body-Based Production of the Influenza Virus Neuraminidase in Escherichia coli. Biomolecules 2022, 12:331.

42. Einsfeldt K, Severo Júnior JB, Corrêa Argondizzo AP, Medeiros MA, Alves TLM, Almeida RV, Larentis AL: Cloning and expression of protease ClpP from Streptococcus pneumoniae in Escherichia coli: Study of the influence of kanamycin and IPTG concentration on cell growth, recombinant protein production and plasmid stability. Vaccine 2011, 29:7136–7143.

43. Masuo S, Saga C, Usui K, Sasakura Y, Kawasaki Y, Takaya N: Glucose-Derived Raspberry Ketone Produced via Engineered Escherichia coli Metabolism. Front Bioeng Biotechnol 2022, 10:843843.

44. Kurokawa M, Ying BW: Precise, High-throughput Analysis of Bacterial Growth. J Vis Exp 2017.

45. Tsuchiya K, Cao YY, Kurokawa M, Ashino K, Yomo T, Ying BW: A decay effect of the growth rate associated with genome reduction in Escherichia coli. BMC Microbiol 2018, 18:101.

46. Liu L, Kurokawa M, Nagai M, Seno S, Ying BW: Correlated chromosomal periodicities according to the growth rate and gene expression. Sci Rep 2020, 10:15531.

47. Ying BW, Matsumoto Y, Kitahara K, Suzuki S, Ono N, Furusawa C, Kishimoto T, Yomo T: Bacterial transcriptome reorganization in thermal adaptive evolution. BMC Genomics 2015, 16:802.

48. Ying BW, Seno S, Kaneko F, Matsuda H, Yomo T: Multilevel comparative analysis of the contributions of genome reduction and heat shock to the Escherichia coli transcriptome. BMC Genomics 2013, 14:25.

49. Ihaka R, Gentleman R: R: A Language for Data Analysis and Graphics. Journal of Computational and Graphical Statistics 1996, 5:299–314.

50. Langmead B, Salzberg SL: Fast gapped-read alignment with Bowtie 2. Nature Methods 2012, 9:357–359.

51. Love MI, Huber W, Anders S: Moderated estimation of fold change and dispersion for RNA-seq data with DESeq2. Genome Biology 2014, 15:550.

52. Storey JD: A direct approach to false discovery rates. Journal of the Royal Statistical Society: Series B (Statistical Methodology) 2002, 64:479–498.

53. Ying BW, Yama K: Gene Expression Order Attributed to Genome Reduction and the Steady Cellular State in Escherichia coli. Front Microbiol 2018, 9:2255.

54. Kanehisa M, Sato Y, Kawashima M, Furumichi M, Tanabe M: KEGG as a reference resource for gene and protein annotation. Nucleic Acids Res 2016, 44:D457–462.

55. Kanehisa M, Furumichi M, Tanabe M, Sato Y, Morishima K: KEGG: new perspectives on genomes, pathways, diseases and drugs. Nucleic Acids Res 2017, 45:D353–D361.

56. Yu G, Wang LG, Han Y, He QY: clusterProfiler: an R package for comparing biological themes among gene clusters. OMICS 2012, 16:284–287.

