## Supplementary figures and tables for "Machine learning-assisted medium optimization revealed the discriminated strategies for improved production of the foreign and native metabolites"

**Supplementary figures S1~S5** **p2~6**
**Supplementary tables S1~S3** **p7**

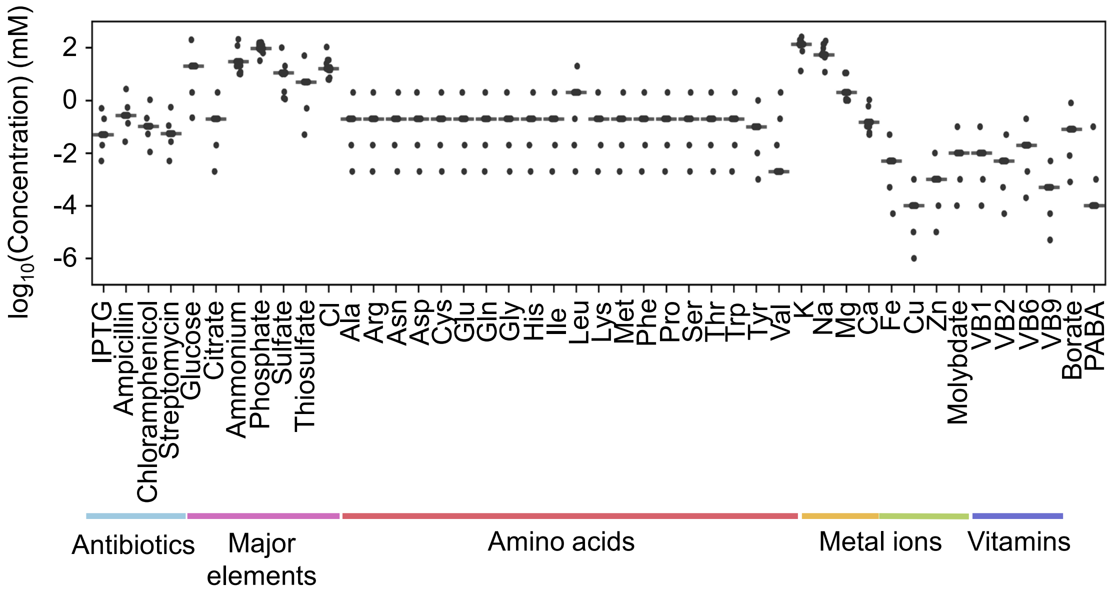


**Figure S1 Tested medium combinations for 4APhe production.** The concentration variation of 48 components comprising 192 medium combinations is shown. The concentrations are indicated on a logarithmic scale. Color variation indicates the categories of elements.

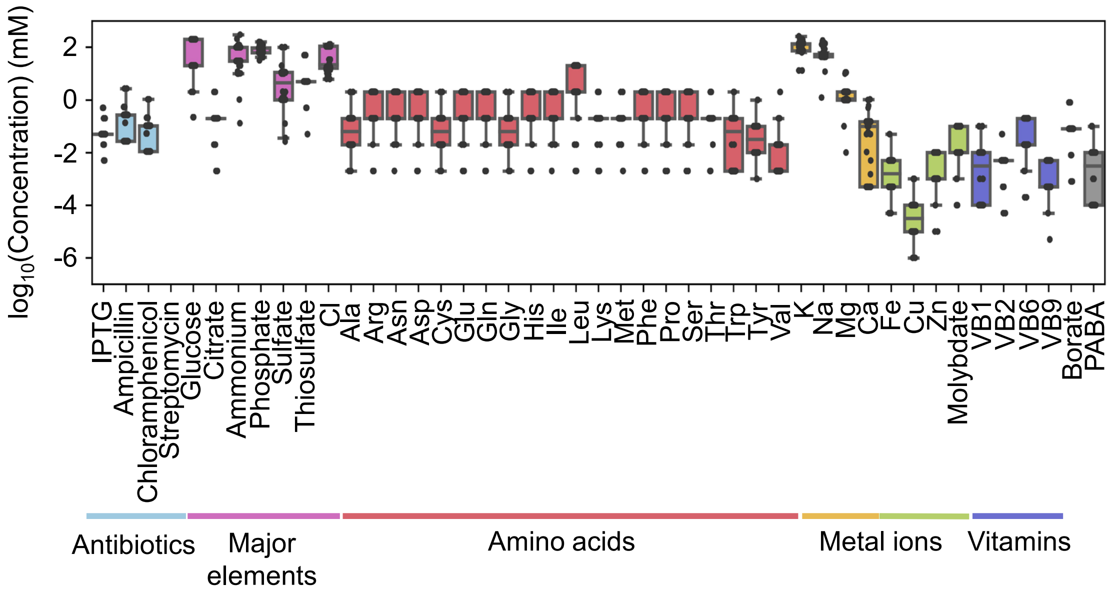


**Figure S2 Tested medium combinations for Tyr production.** The concentration variation of 48 components comprising 378 medium combinations is shown. The concentrations are indicated on a logarithmic scale. Color variation indicates the categories of elements.

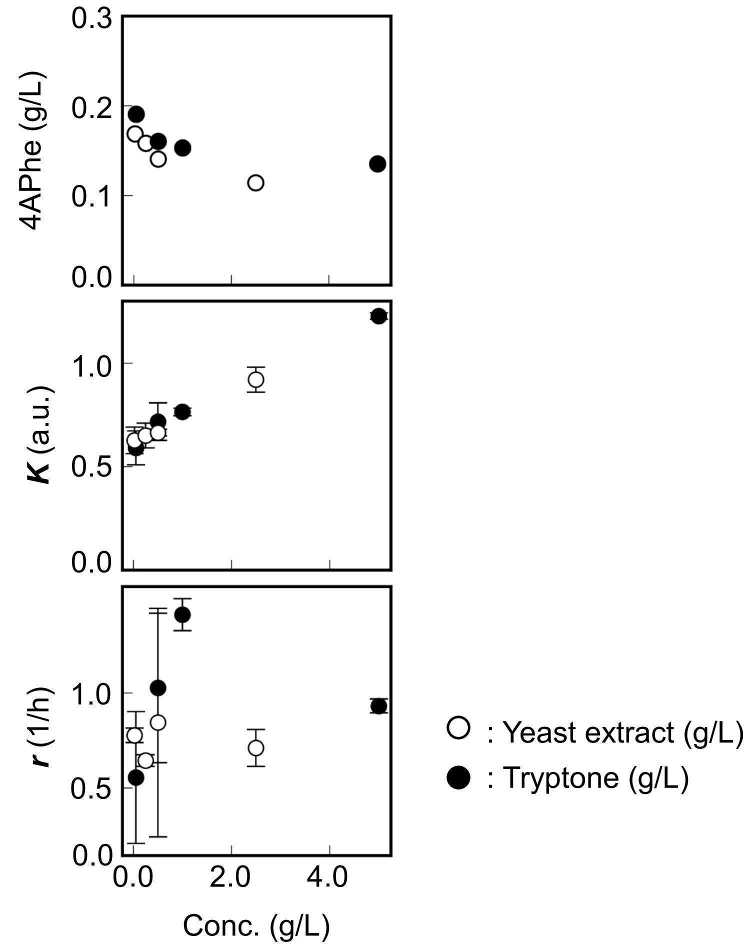


**Figure S3 Contribution of nutritional richness to bacterial growth and productivity.** The *E. coli* strain holding the synthetic pathways for producing 4APhe was cultured in the media supplemented with the natural gradients of yeast extract or tryptone, represented by the open and closed circles, respectively. *K* and *r* represent the maximal density and growth rate, respectively. Standard errors of experimental replications are indicated.

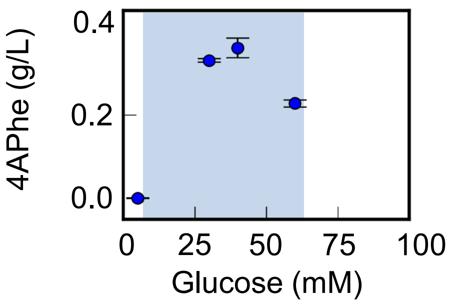


**Figure S4 4APhe production in response to the concentration gradient of glucose.** The *E. coli* strain holding the synthetic pathways for 4APhe production was cultured in 5 ml of media. The shadow represents the concentration gradient of glucose recommended by the CART model. Standard errors of experimental replications are indicated.

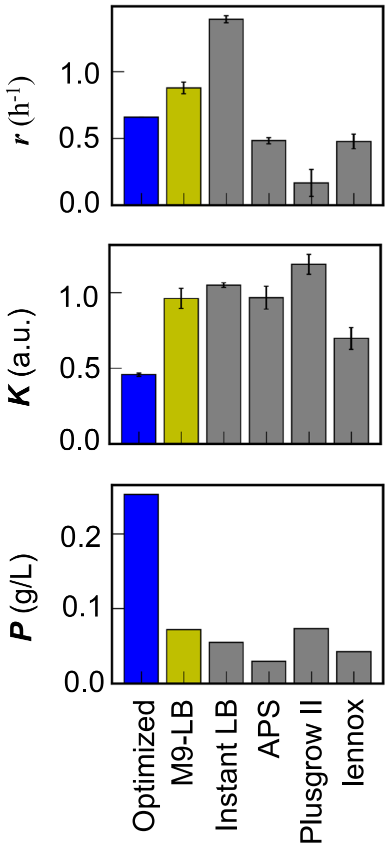


**Figure S5 Comparison of the fine-tuned medium to the commercially available media.** The *E. coli* strain holding the synthetic pathways for producing 4APhe was cultured in the optimized medium, M9-LB mixed medium, and four commercially purchased media, shown in blue, yellow, and grey, respectively. *K*, *r*, and *P* represent the maximal density, growth rate, and 4APhe production (yield). Standard errors of experimental replications are indicated.

**Table S1 Dataset of medium combinations to the growth and production of the strain producing 4APhe.** The logarithmic concentrations are shown, in which zero concentration was replaced with the logarithmic value of one-hundredth of its minimal concentration used to prepare the medium combinations.

(Datasheet elsewhere)

**Table S2 Dataset of medium combinations to the growth and production of the strain producing Tyr.** The logarithmic concentrations are shown, in which zero concentration was replaced with the logarithmic value of one-hundredth of its minimal concentration used to prepare the medium combinations.

(Datasheet elsewhere)

**Table S3 The number of components contributing to 70% of feature importance.** $K$, $r$, and $P$ represent the maximal density, growth rate, and production (yield). The values (%) in the brackets are the sum of the feature importance attributed to the components indicated.

| | 4APhe | Tyr |
|---|---|---|
| $r$ | 6 (70%) | 4 (73%) |
| $K$ | 5 (72%) | 6 (70%) |
| $P$ | 7 (71%) | 14 (70%) |
